# Synthetic derivatives of vinpocetine as antiproliferative agents

**DOI:** 10.1101/2025.03.15.643482

**Authors:** Mihira Gutti, Melanie Tsui, Stella Yang, Selina Xi, Jennifer Luo, Arshia Desarkar, Yining Xie, Mirabelle Feng, Udbhav Avadhani, Shloka Raghavan, Elena Brierley-Green, Erika Yu, Edward Njoo

## Abstract

Vincamine is an indole alkaloid initially isolated from plants of the *Vinca* genus and has previously been demonstrated to have antioxidant, hypoglycemic, and hypolipidemic activities. Vinpocetine, a synthetic derivative of vincamine with an enhanced pharmacological profile, has demonstrated promising antiproliferative properties. While previously reported vinpocetine derivatives have undergone extensive investigation for their pharmacological properties, the role of the E-ring ethyl ester in the anticancer properties of compounds with this scaffold has not yet been fully described. Here, we report the antiproliferative activity of two vinpocetine analogs with modifications at the E-ring. MTT assays revealed that reduction of the ethyl ester to an alcohol exhibited strong dose-dependent antiproliferative activity across five mammalian cell lines, but did not induce significant markers of apoptosis or necrotic death as determined by FITC/Annexin V and cell cycle flow cytometry, respectively. We further observe that vinpocetine and both analogs exhibit dose-dependent modulation of a TCF/LEF reporter cell, which appears to be decoupled from trends in antiproliferative activity Altogether, this work demonstrates the potential for E-ring modifications of vinpocetine as anticancer agents.

## Introduction

Vinca alkaloids and their derivatives, which share biosynthetic origins from tryptamine and related aminoindoles (Zhu et al., 2015), have demonstrated remarkable potency for diverse applications in chemical biology as neuroactive agents (Hussain et al., 2018), anti-inflammatory agents (Jeon et al., 2010), immunomodulators (Byrd & Isa, 1981), and as lead compounds for the treatment of cancer (Jordan, Thrower & Wilson, 1991; Martino et al., 2018). To date, three such alkaloids have received FDA approval as chemotherapy agents, such as vincristine and vinblastine, which were first to be FDA-approved in 1961 and 1963, respectively (Carbone et al., 1963; Sears & Boger 2015) **(Fig. 1)**.

**Figure 1.**
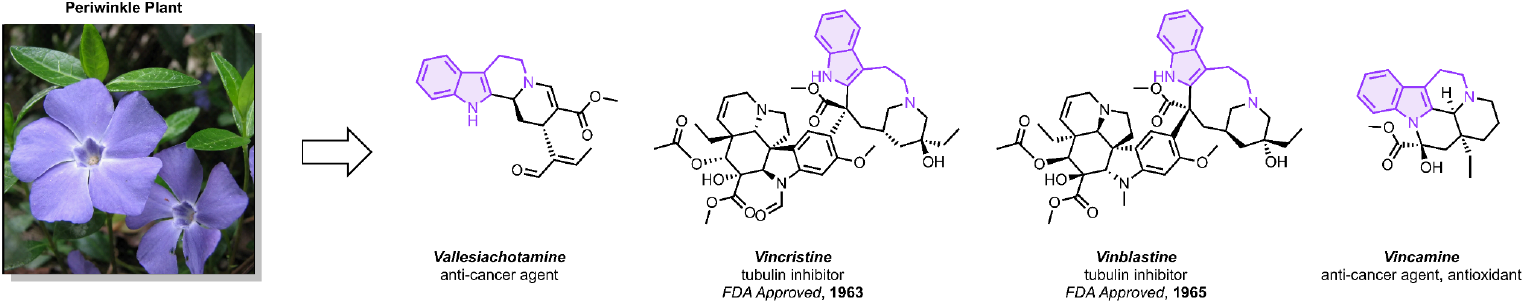
Vinca alkaloids show diverse biological activity, including anticancer activity. Photo by and © 2007 Jina Lee.

Vincamine is an indole alkaloid initially isolated from the periwinkle plant *Vinca minor*, later identified in *Vinca rosea* (*Catharanthus roseus*), and has previously demonstrated to have antioxidant, hypoglycemic, and hypolipidemic activities (Wu, Ye & Zhang, 2018; Nandini & Naik, 2019; Ren et al., 2023). Additionally, vincamine has also been studied in preclinical models for the treatment of type 2 diabetes mellitus and drug addiction (Wang et al., 2020; Norwood et al., 2020). Due to its potent antioxidant effects, vincamine has been further investigated for the treatment of cerebrovascular diseases (Vas & Gulyás, 2005; Nemes, 2010; Abdel-Salem et al., 2016; Shalaby et al., 2019; Ren et al., 2023).

A synthetic derivative of vincamine, vinpocetine, was developed to enhance its pharmacological profile and has been in clinical use in several European countries including Hungary, Germany, and Poland as well as Japan for the treatment of cerebrovascular diseases for over 30 years (Willson, 2009; Zhang, Li & Yan, 2018). In recent years, studies have found vinpocetine to have a variety of pharmacological activities, including anti-inflammatory, cardioprotective, and hepatoprotective activity (Dubey et al., 2020). Remarkably, both vincamine and vinpocetine have also demonstrated promising antiproliferative and anticarcinogenic activities, highlighting their broader therapeutic potential (Huang et al., 2012; Al-Rashed et al., 2021; Ciorîţă et al., 2021; Zhang et al., 2021; Dhyani et al., 2022; Ren et al., 2023; Mohammed et al., 2024). Various biological targets have been proposed as putative targets through which vincamine, vinpocetine, and their derivatives elicit their biological effects, including phosphodiesterase 1 (PDE1) (Shekarian et al., 2020; Sharma et al., 2022), Wnt1 (Ji et al., 2023), and nuclear factor-κB (Nf-κB) (Jeon et al., 2010).

Previously reported vinpocetine derivatives have undergone extensive investigation for their pharmacological properties (Dong et al., 2023; Dong et al., 2024). Specifically, synthetic derivatization of the E-ring, reported with initial reduction followed by carbamate protection (Pan et al., 2020) and hydrolysis followed by the introduction of an amide (Sheng et al., 2011), aldehyde (Nemes et al., 2008), nitroxy group (Kawashima et al., 1993), or fluorine atom (Nag et al., 2019), has been studied for its PDE1 inhibition, antioxidant effects, artery blood flow effects, and blood-brain barrier permeability (Dong et al., 2024). However, the anticancer properties of vinpocetine derivatives with E-ring modifications have yet to be fully described. Reduction of the E-ring ester or hydrolysis to the corresponding acid are two immediate strategies for changes to the vinpocetine scaffold **(Scheme 1)**, and in this study, we specifically sought to identify the effect of these two subtle chemical modifications on the biological activity of vinpocetine and its analogs in mammalian cancer models. While the corresponding alcohol and acid have been previously explored for their PDE1A inhibition and antioxidant activity (Sheng et al. 2011; Pan et al., 2020), their potential in applications related to cancer has not been fully explored.

**Scheme 1.**
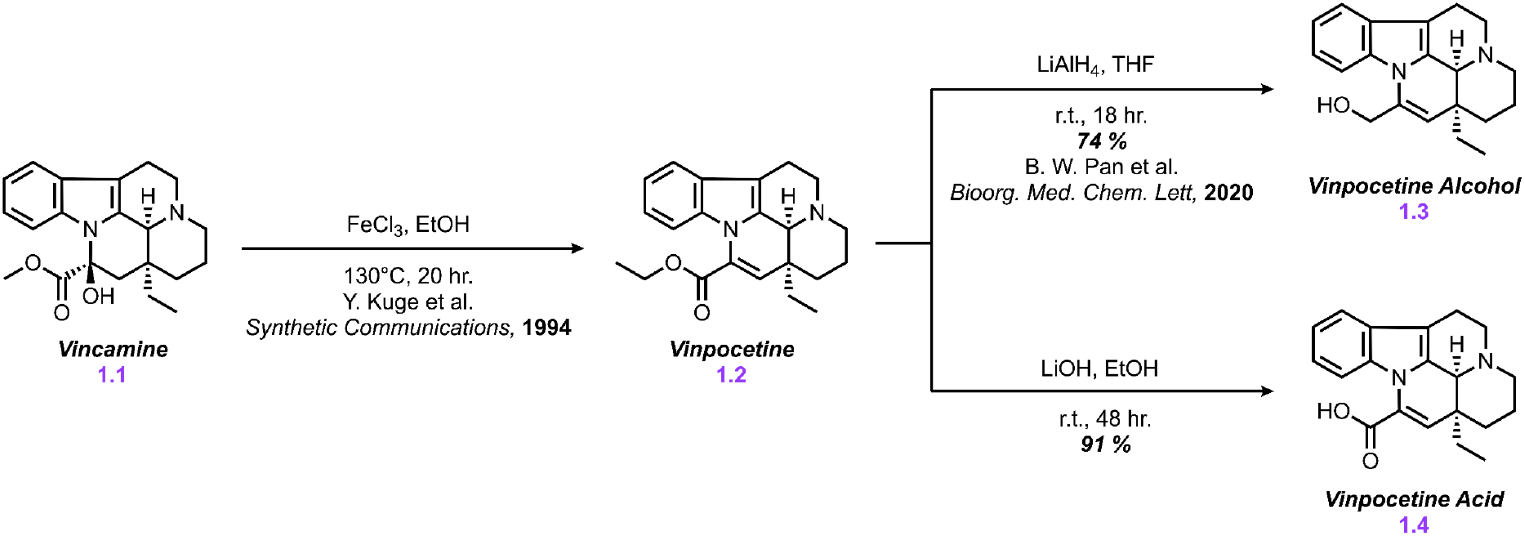
Synthesis of Vinpocetine derivatives.

## Materials & Methods

### General Information

All reagents, solvents, and analytical standards were purchased from AK Scientific and Sigma Aldrich and were used without further purification unless otherwise stated. McCoy’s 5A Media, Dulbecco’s Modified Eagle’s Medium (DMEM), RPMI-1640 Medium, penicillin-streptomycin, and 0.25% trypsin-EDTA were all obtained from Tribioscience (Sunnyvale, CA). Fetal bovine serum (FBS) was obtained from Gibco. MTT and synthetic reagents were obtained from AK Scientific.

### Cell Culture

Human colorectal cancer (HCT-116 and HT-29) and human non-small-cell lung cancer (HTB-54) cell lines were obtained from the European Collection of Authenticated Cell Cultures (ECACC) and maintained in McCoy’s 5A Media (Tribioscience; Sunnyvale, CA) supplemented with 10% fetal bovine serum (FBS) from Gibco and 1% penicillin-streptomycin from Tribioscience (Sunnyvale, CA). Human embryonic kidney (HEK-293TN) cell lines were obtained from System Biosciences (Palo Alto, CA) and donated by SunnyBay Biotech (Fremont, CA), and cultured in Dulbecco’s Modified Eagle’s Medium (DMEM) from Tribioscience, supplemented with 10% fetal bovine serum (FBS) and 1% penicillin-streptomycin. Human breast cancer (MDA-MB-231) cell lines, donated by SunnyBay Biotech (Fremont, CA), were cultured in RPMI-1640 Medium from Corning supplemented with 10% fetal bovine serum (FBS) and 1% penicillin-streptomycin. Leading Light® Wnt Reporter 3T3 mouse embryonic fibroblast cells from Enzo Life Sciences (Cat. # ENZ-61002-0001) were maintained in DMEM, supplemented with 10% fetal bovine serum (FBS) and 1% penicillin-streptomycin. All cells were grown in 25 cm^2^ and 75 cm^2^ flasks (Bio Basics) in a humidified incubator at 37°C in 5% CO_2_.

### Cell Viability Assays

Cells (HCT-116, HT-29, HEK-293TN, MDA-MB-231, HTB-54) were seeded in a 96-well tissue culture-treated flat bottom plate at 20-25% confluency and incubated in a humidified incubator at 37°C in 5% CO_2_. Treatment was conducted at 48 and 72 hours at different concentrations of compound in DMSO (500 μM, 250 μM, 50, 25 μM, 5 μM, 2.5 μM, 500 nM, 200 nM). After incubation, 10 μL of a 5 mg/mL MTT (3-(4,5-dimethylthiazol-2-yl)-2,5-diphenyltetrazolium bromide) solution in 1X PBS was added to each well, and then incubated for 2 hours at 37°C. Then, each well was replaced with 100 μL of DMSO before measuring absorbance at 570 nm using a Molecular Devices SpectraMax 250 Microplate Spectrophotometer. Percent cell viability was determined through the ratio of absorbance of treated cells to the negative control cells (DMSO, without compound). IC_50_ values were calculated on GraphPad Prism 10.

### Cell Cycle Flow Cytometry

Cells (HCT-116, HEK-293TN, MDA-MB-231) were seeded at 70% confluency in a 6-well tissue culture-treated flat bottom plate and incubated in a humidified incubator at 37°C in 5% CO_2_ for 24 hours, after which treatment was conducted at the highest concentration of 500 µM. After incubation for a subsequent 24 hours, supernatant media was aspirated off, and the cells were trypsinized with 500 µL 0.25% trypsin-EDTA. Trypsin was then deactivated with 500 µL of cell-line respective media (McCoy’s for HCT-116, DMEM for HEK-293TN, RPMI-1640 for MDA-MB-231), and cells with supernatant media were centrifuged for 5 minutes at 1.9 rpm until cell pellet formed. Supernatant media was removed, and the cell pellet was resuspended in 100 µL of 1X PBS. After centrifuging for an additional 4 minutes, PBS was removed and cells were resuspended in cold 70% ethanol. Cells were left to fix on ice for 30 minutes, after which they were centrifuged into pellets for an additional 4 minutes, ethanol was removed, and cells were resuspended in 200 µL PBS. 50 µL RNAse (1 mg/mL) was added to the cells, which were subsequently incubated for 5 minutes at 37°C in 5% CO_2_. After incubation, the cells were stained with 1 µL Propidium Iodide stain for 30 minutes prior to analysis by flow cytometry. Cell cycle analysis was run on a BD C6 Accuri Flow Cytometer to 50,000-100,000 events for each compound. Analysis of results was performed through BD C6 Accuri Software. Single cells were gated to exclude debris and cell aggregates based on forward scatter and side scatter. Based on PI/count gating, the percentage of cells that were in the G0/G1 phase, S phase, and G2/M phase were calculated for each sample.

### Apoptosis Flow Cytometry

Cells (HCT-116, MDA-MB-231) were seeded at 70% confluency in a 12-well tissue culture-treated flat bottom plate and incubated in a humidified incubator at 37°C in 5% CO_2_. Treatment was conducted for 72 hours at the highest concentration of 500 µM. After incubation, the supernatant media was aspirated off, and the cells were trypsinized with 400 µL 0.25% trypsin-EDTA. Subsequently, the trypsin was deactivated with 400 µL of cell-line respective media (McCoy’s for HCT-116, RPMI-1640 for MDA-MB-231), and cells with supernatant media were centrifuged at 1.9 rpm until cell pellet formed. The cell pellet was then washed with 100 µL of 1X PBS, re-pelleted by centrifugation, and stained with 100 µL 1X Annexin binding buffer (Invitrogen), 5 µL FITC-Annexin stain (Southern Biotech), and 1 µL 1.0 mg/mL propidium iodide (PI) stain on ice for 15 minutes. Apoptosis analysis was performed by flow cytometry on a BD C6 Accuri Flow Cytometer to 25,000 events for each sample. Single cells were gated to exclude debris and cell aggregates based on forward scatter and side scatter. Based on Annexin V/PI quadrant gating, the percentage of cells that were viable in the early apoptotic, late-stage apoptotic, and necrotic populations were quantified and expressed as percentages of total cell populations for each sample.

### Wnt1 Luminescence-based Reporter Assay

Leading Light® Wnt Reporter 3T3 mouse embryonic fibroblast cells were obtained from Enzo Life Sciences (cat. ENZ-61002-0001) and seeded at 80% confluency in 96-well flat bottom plates (Corning Costar). After 24 hours of incubation at 37 °C with 5% CO_2_, the plates were treated for 24 hours with **1.1 - 1.4** at 500 μM, 250 μM, 50 μM, and 25 μM and with and without CHIR-99021 (AK Scientific), a known GSK-3β inhibitor at a final concentration of 10 μM. A negative control of 0.5% v/v DMSO and a background control were included. After 24 hours of incubation at 37°C with 5% CO_2_, 70 μL of 3X Firefly Assay Buffer (15 mM Dithiothreitol, 0.45 mM ATP, 4.2 mg/mL D-luciferin, Triton X-100 Lysis Buffer (0.1082 M Tris-HCl powder, 0.0419 M Tris-base powder, 75 mM NaCl, 3 mM MgCl2, 0.25% Triton X-100, H_2_O)) were added to each well on top of the cell media. Immediately after, luminescence was quantified using a Tecan SpectraFLUOR Plus plate reader.

## Results and Discussion

### Chemical Synthesis

Given the critical role of the E-ring in the biological activity of vinpocetine, particularly the influence of the ethyl ester, we synthesized two analogs to investigate the effect of structural modifications on its activity. Compound **1.3** was reported through lithium aluminum hydride reduction in tetrahydrofuran in 74% isolated yield (Pan et al., 2020) **(Scheme 1)**. Additionally, compound **1.4** was synthesized by hydrolysis of the ethyl ester of vinpocetine to the corresponding acid with lithium hydroxide in ethanol in 91% isolated yield **(Scheme 1)**.

### MTT Assays

In order to assess the antiproliferative activitiy of compounds **1.1 - 1.4**, we performed MTT assays in five mammalian cell lines, including HCT-116 and HT-29 human colorectal carcinoma, MDA-MB-231 human breast cancer, HTB-54 human non-small cell lung cancer, and HEK-293TN human embryonic kidney cells **(Fig. 2)**.

**Figure 2.**
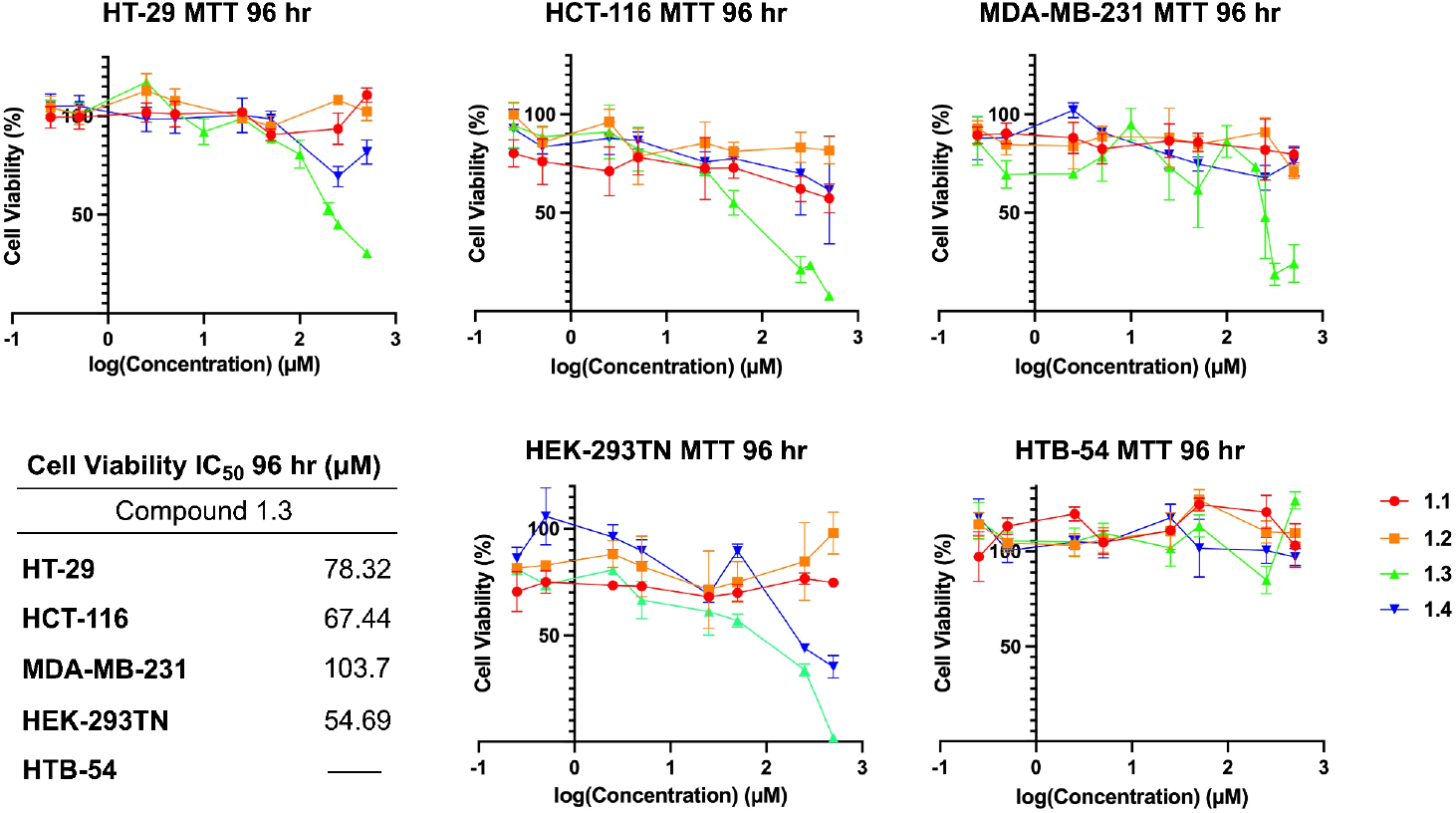
Cell viabilities of compounds **1.1 - 1.4** in HT-29, HCT-116, MDA-MB-231 breast cancer cells, HEK-293TN human embryonic kidney cells, and HTB-54 non-small cell-lung cancer at a 96-hour time point and IC_50_ values of compound **1.3** across cancerous cell lines.

After 96 hours, compound **1.3** exhibited dose-dependent antiproliferative activity, exhibiting an IC_50_ value in the double-digit to triple-digit micromolar range across all cell lines except HTB-54 cells. Comparatively, while vincamine **1.1**, vinpocetine **1.2**, and acid **1.4** exhibited minimal antiproliferative activity, alcohol **1.3** had IC_50_ values of 78.32 μM, 67.44 μM, 103.7 μM, and 54.69 μM in HT-29, HCT-116, MDA-MB-231, and HEK-293TN cells, respectively. The increase in potency in the cells treated with compound **1.3** suggests that the activity of vinpocetine derivatives is dependent on modifications on the E-ring.

### Cell Cycle Analysis by Flow Cytometry

In order to analyze the cell cycle arrest capabilities of compounds **1.1 - 1.4**, we performed cell cycle flow cytometry assays in three mammalian cell lines, including HCT-116 human colorectal carcinoma cells, MDA-MB-231 human breast cancer cells, and HEK-293TN human embryonic kidney cells **(Fig. 3)**. We observe minimal cell-cycle arrest via cell-cycle analysis in HCT-116 cells in the S-phase with alcohol **1.3**, but no significant effect with other compounds, suggesting that the primary mechanism through which **1.3** exhibits its antiproliferative activity is not related to mechanical inhibition of the cell cycle, unlike vincristine and vinblastine, which function through mitotic inhibition.

**Figure 3.**
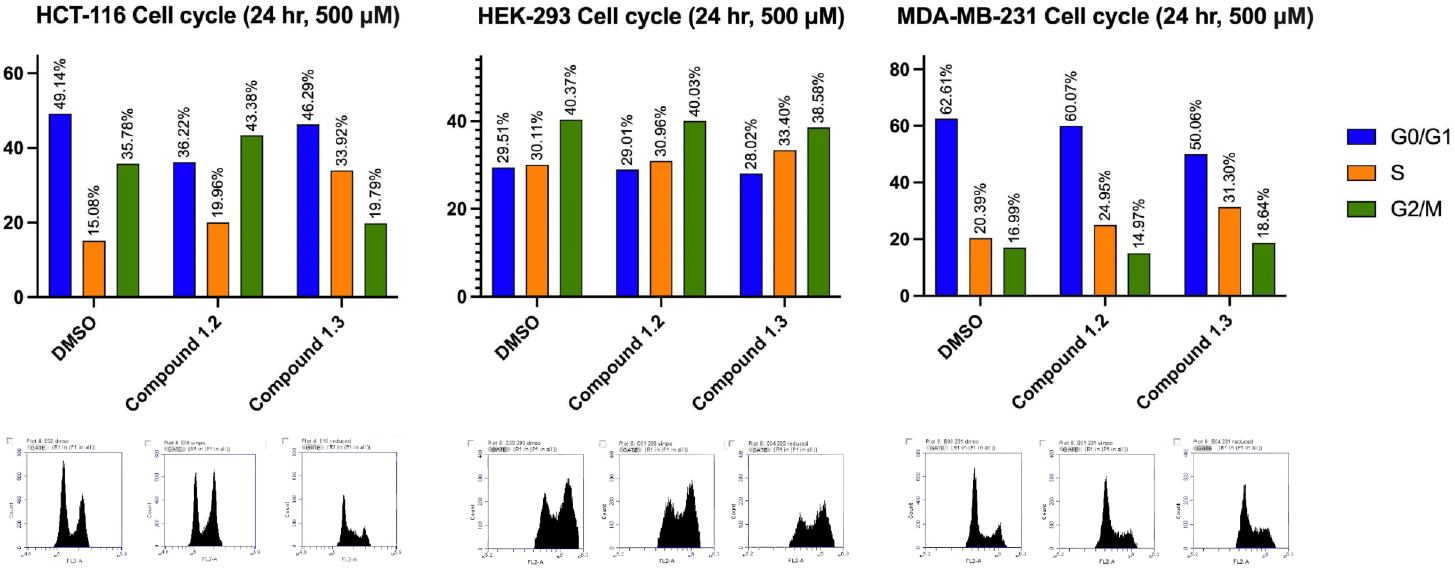
Flow cytometry of HEK-293TN, MDA-MB-231, and HCT-116 cells treated with compounds **1.2 - 1.3**. The cell cycle distribution of each treated population was determined via flow cytometry using propidium iodide (PI) staining after ethanol fixation to determine the cell-cycle arrest in cells treated with compounds **1.2 - 1.3** and a DMSO negative control over a 24-hour period.

### Apoptosis Assay by Flow Cytometry

Next, we sought to evaluate whether our compounds potentially functioned through a pro-apoptotic pathway. To detect and quantify the level of apoptosis of compounds **1.1 - 1.4**, we performed apoptosis flow cytometry assays in HCT-116 and MDA-MB-231 human colorectal carcinoma cells **(Fig. 4)**, utilizing a propidium iodide (PI) and FITC-labeled Annexin V double stain. We found that these compounds do not appear to function via a pro-apoptotic pathway due to lack of FITC signals. We also observed minimal PI signals in both cell lines. This suggests that while **1.3** exhibits strong antiproliferative activity, this attenuation of cell proliferation results in minimal differences in cytotoxicity.

**Figure 4.**
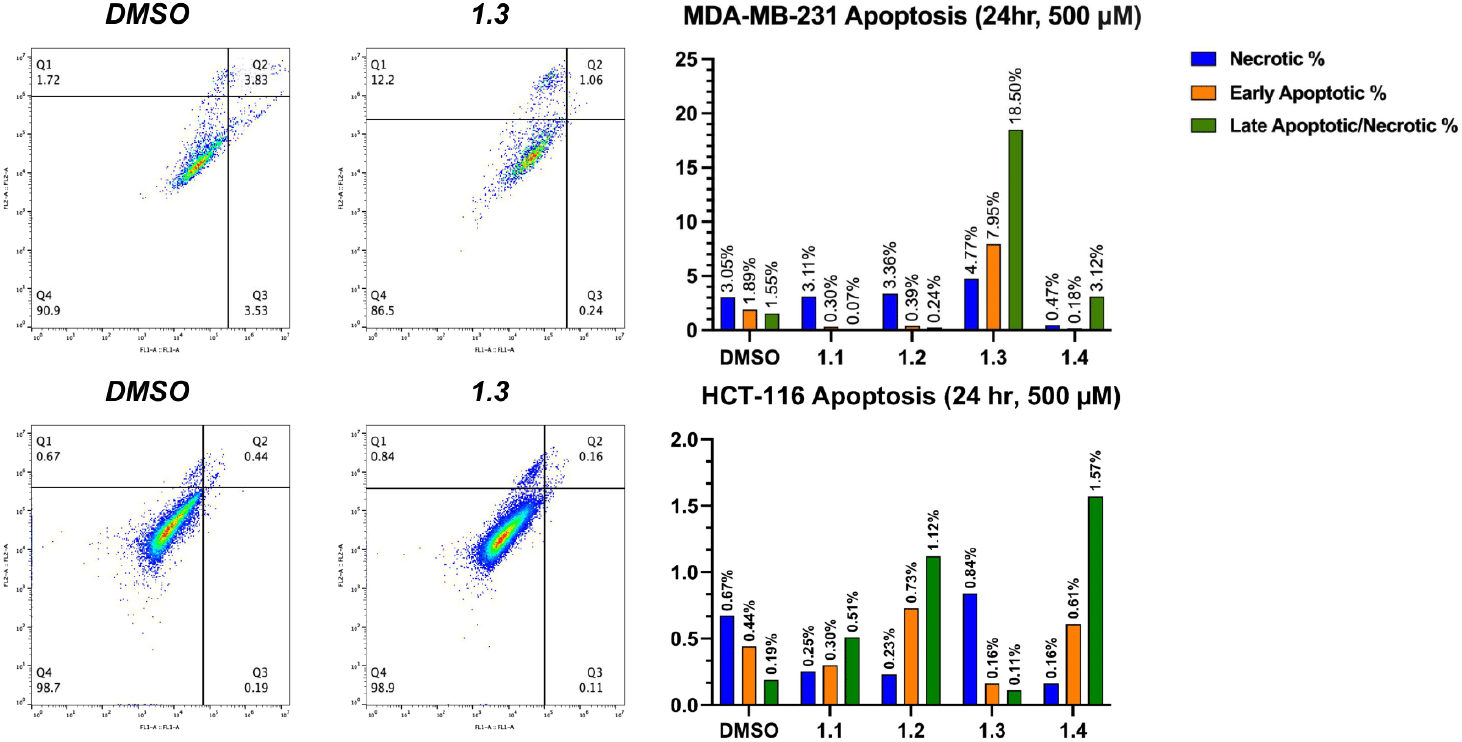
Flow cytometry of MDA-MB-231 and HCT-116 cells treated with compounds **1.1 - 1.4**. Standard Annexin-V/Propidium Iodide protocol was followed to determine the percentage of cells undergoing apoptosis in cells treated with compounds **1.1 - 1.4** and a DMSO negative control over a 48-hour period.

### Wnt1 Luminescence-based Reporter Assay

To determine the effect of compounds **1.1 - 1.4** on Wnt1 signaling, we employed an *in vitro* reporter cell luciferase assay using a stable transfected Wnt1-responsive 3T3 murine fibroblast cell line expressing a TCF/LEF-activated firefly luciferase gene. Compounds **1.1 - 1.4** were administered at 500 µM, 250 µM, 50 µM, 25 µM concentrations and incubated for 24 hours, after which a luciferin substrate mix was added, and luminescence was observed on a 96-well plate reader. In the absence of a GSK-3β kinase inhibitor, vinpocetine **1.2** and its analogs **1.3** and **1.4** exhibit dose-dependent inhibition of fLuc activity at the highest concentrations, while vincamine 1.1 is largely inactive in this assay **(Fig. 5)**. However, when co-delivered with 10 μM CHIR-99021, a selective GSK-3β kinase inhibitor (Cohen & Goedert, 2004), compounds **1.2 - 1.4** appear to have a net inhibitory effect by fLuc signal at high concentrations but a net activation at concentrations above 50μM. Additionally, while vinpocetine **1.2** and its carboxylic acid **1.4** exhibit minimal antiproliferative activity, similar trends are observed in our luciferase assay, suggesting that while compounds of this class may alter Wnt1 signaling in cells, this is not the basis for the antiproliferative activity exhibited by alcohol **1.3**.

**Figure 5.**
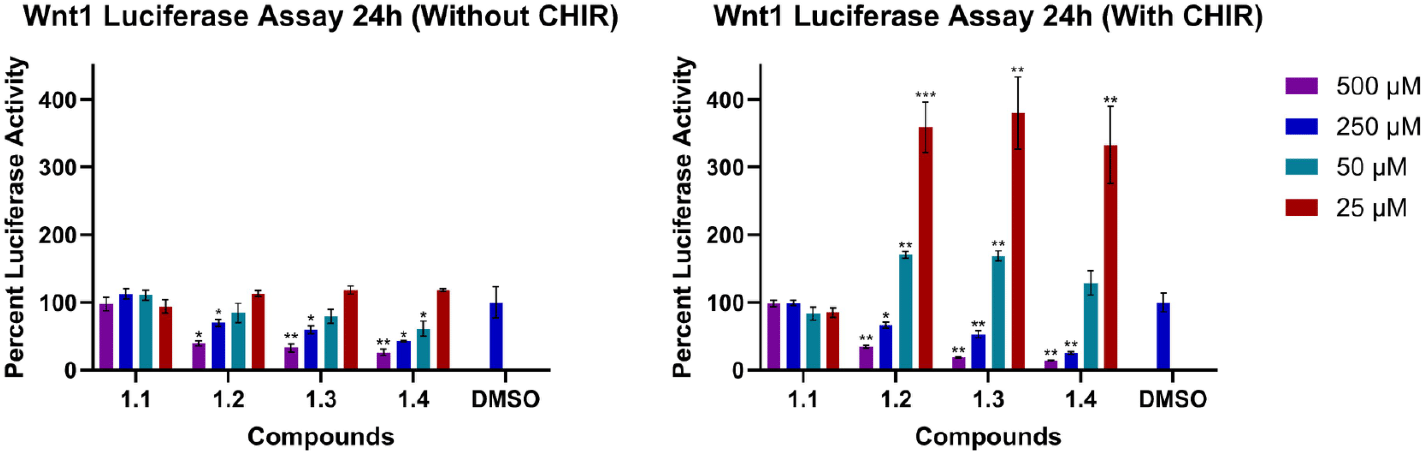
Activity of Compounds **1.1 - 1.4** in a Wnt1-dependent luciferase reporter cell assay. Luciferase reporter activity from Leading Light® Wnt Reporter 3T3 mouse fibroblast cells was quantified through readout of the resulting luminescence from the oxidation of the luciferin substrate. A negative control was established with 0.5% v/v DMSO, and data is represented as means relative to a background control ± SD compared with the negative control using a Welch’s t-test (n = 4) (*P<0.05, **P<0.01, ***P<0.001).

## Conclusions

Inspired by the diverse biological activities of several *Vinca* alkaloids, we sought to identify the effects of simple modifications of the E-ring ethyl ester of vinpocetine in a panel of mammalian cancer cell lines. Here, we prepared two E-ring analogs of vinpocetine with a primary alcohol **1.3** and carboxylic acid **1.4** and evaluated their potencies in antiproliferative activity in HCT-116 and HT-29 human colorectal cancer cells, MDA-MB-231 human breast cancer cells, HTB-54 human non-small cell lung cancer cells, and HEK-293TN human embryonic kidney cells. Consistent with prior reports on the limited toxicity of vinpocetine, we observed that both this and its parent natural product, vincamine, exhibited minimal effects on the five cell lines in this study. Similarly, acid **1.4** exhibited slight antiproliferative activity only at the highest concentrations (500 μM). However, alcohol **1.3** exhibited strong dose-dependent antiproliferative activity uniformly across all five mammalian cell lines with IC_50_ values ranging from 54.69 μM in HEK-293TN to 103.7 μM in MDA-MB-231 μM.

Further mechanistic evaluation performed by flow cytometry suggests that alcohol **1.3** exhibits minimal cell cycle arrest, suggesting that mechanical mitotic inhibition is not involved in the primary mechanism through which **1.3** acts. Additional flow cytometry experiments revealed that cells treated with alcohol **1.3** do not significantly produce positive signals in accumulated Annexin V, suggesting that the observed antiproliferative activity of **1.3** is not attributed to the activation of pro-apoptotic pathways. Finally, the differential activity exhibited by the compounds in this study in a Wnt1/TCF/LEF fLuc reporter assay suggests that the mechanism of action through which **1.3** exhibits antiproliferative activity is not likely to be through direct involvement of the Wnt1 pathway.

In conclusion, we show that, while vinpocetine and vincamine exhibit very limited antiproliferative activity in model cancer cell lines, modifications of the E-ring ethyl ester may augment the biological potency of vinpocetine analogs as anticancer agents. Specifically, a reduced vinpocetine analog bearing an E-ring allylic alcohol **1.3** exhibits dose-dependent antiproliferative activity in five mammalian cell lines. The results of this work indicate that E-ring synthetic derivatization can be a strategy for the development of future vinpocetine analogs with anticancer activity.

## Acknowledgments

The authors express their gratitude to Dr. Feng Wang-Johanning and Dr. Gary Johanning from SunnyBay Biotech for supplying the HEK-293TN and MDA-MB-231 cells. They also thank Dr. Stephen Lynch for access to high-field NMR spectra at the Stanford University NMR facility.

## Notes

### Competing Interest Statement

The authors have declared no competing interest.

