## Supporting Information Document for "Synthetic derivatives of vinpocetine as antiproliferative agents"

### ***Synthetic derivatives of Vinpocetine as anti-proliferative agents for the treatment of solid tumors***

CONTENTS

A. General Information SI-1
   a. Materials SI-1
   b. Equipment SI-1

B. Experimental Procedures SI-2
   a. Cell Culture SI-2
   b. Cell Viability Assays SI-2
   c. Cell Cycle Flow Cytometry SI-2
   d. Apoptosis Flow Cytometry SI-3
   e. Wnt-1 Luminescence-based Reporter Assay SI-3

C. Chemical Synthesis SI-4
   a. 1.3 Vinpocetine Alcohol SI-4
   b. 1.4 Vinpocetine Acid SI-8

D. Supplemental Tables SI-12

### A. General Information

#### a. Materials

Solvents and reagents used in all reactions and purification processes were ACS grade or higher, were used without additional purification, and were purchased from Sigma Aldrich or AK Scientific. Deuterated solvents were purchased from Cambridge Isotope Laboratories, Acros Organics, or Martek Isotopes and were used without further purification. All other reagents, catalysts, and chemicals were purchased from commercial sources and used without further purification unless otherwise stated.

#### b. Equipment

$^{1}H$ and $^{13}C\{1H\}$ NMR spectra were acquired on a Varian INOVA 400 MHz nuclear magnetic resonance spectrometer, a Bruker Avance Neo 400 MHz nuclear magnetic resonance spectrometer, a JEOL 500 MHz nuclear magnetic resonance spectrometer, or a Nanalysis NMReady 60Pro 60 MHz multinuclear benchtop nuclear magnetic resonance spectrometer and were processed on the MestreNova software package. and were processed on the MestreNova software package. $^{1}H$ and $^{13}C$ NMR spectra are reported in parts per million (ppm) relative to the residual solvent peak (Methanol-*d*4 = 3.34 ppm) as follows: chemical shift (δ), multiplicity (app = apparent, b = broad, s = singlet, d = doublet, t = triplet, q = quartet, m = multiplet, or combinations thereof), coupling constant(s) in Hz, integration. $^{13}C$ chemical shifts are reported relative to the residual solvent peak (Methanol-*d*4 = 48.3 ppm). Mass spectra were obtained on a Waters Micromass Quattro triple quadrupole mass spectrometer and were processed on the MestreNova software package. Infrared spectra were collected on a Thermo Scientific Nicolet iS5 Fourier transform infrared (FT-IR) spectrometer equipped with a Thermo iD5 attenuated total reflectance (ATR) assembly. Flow cytometry data was collected on a BD C6 Accuri Flow Cytometer and luminescence data was quantified using a Tecan SpectraFLUOR Plus plate reader.

### C. Chemical Synthesis

#### a. Compound 1.3 Vinpocetine Alcohol

**Preparation of Vinpocetine Alcohol (1.3)**

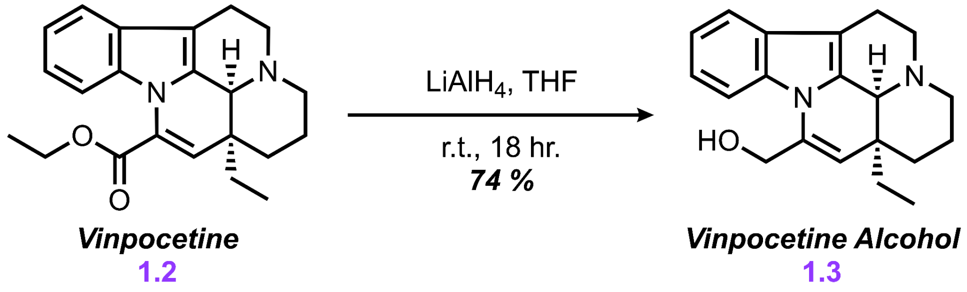


**Chemicals:**
Vinpocetine (AK Scientific, 95%): Used without further purification
Lithium aluminum hydride (Alfa Aesar, 1.0 M in tetrahydrofuran): Used via SureSeal

**Procedure:** To an oven-dried 25 mL 3-neck flask containing vinpocetine (700 mg, 2.0 mmol, 2.0 equiv) suspended in tetrahydrofuran (10 mL) cooled to 0ºC in an ice bath dewar was added lithium aluminum hydride (4.0 mL, 4.0 mmol, 4.0 equiv) via syringe. The resulting suspension was stirred and allowed to reach room temperature overnight. After 18 hr. TLC analysis showed full reduction of ester **1.2** to alcohol **1.3**. The reaction mixture was quenched with EtOAc and was purified by crystallization to obtain a white powder, affording vinpocetine alcohol **1.3** (456 mg, 74% yield).

**Characterization Data of Vinpocetine Alcohol 1.3**

**TLC ($R_f$):** 0.5 (95% EtOH/ 5% N-Heptane), UV active, purple-blue spot by ninhydrin stain

**LCMS (ESI-MS)**: Calculated for $[C_{20}H_{24}N_2ONa]^+[M+Na]^+$: 331.415; found: 331.636

**FT-IR (ATR, $cm^{-1}$)**: 3047.76, 2931.68, 2851.42, 2358.25, 2243.43, 1651.09, 1451.73, 1416.52, 1396.07, 1286.70, 1216.34, 1139.97, 1112.52, 1021.44, 906.04, 808.27, 728.22, 668.20, 646.98, 598.80

**$^1$H NMR** (400 MHz, $CD_3OD$) δ 7.67 (d, *J* = 8.4 Hz, 1H), 7.40 (dd, *J* = 7.7, 1.3 Hz, 1H), 7.13 (ddd, *J* = 8.5, 7.0, 1.4 Hz, 1H), 7.09 – 7.01 (m, 1H), 5.15 (s, 1H), 4.75 (dd, *J* = 13.1, 1.0 Hz, 1H), 4.57 (d, *J* = 13.1 Hz, 1H), 4.17 (d, *J* = 2.6 Hz, 1H), 3.29 (dd, *J* = 13.4, 5.8 Hz, 1H), 3.19 (ddd, *J* = 13.6, 11.5, 5.2 Hz, 1H), 3.02 (dddd, *J* = 17.3, 11.5, 6.1, 2.7 Hz, 1H), 2.72 (td, *J* = 11.8, 2.8 Hz, 1H), 2.62 (dp, *J* = 11.4, 1.9 Hz, 1H), 2.52 (ddd, *J* = 16.0, 4.9, 2.0 Hz, 1H), 2.07 – 1.87 (m, 1H), 1.81 – 1.63 (m, 2H), 1.52 – 1.37 (m, 2H), 1.17 – 1.06 (m, 1H), 1.03 (t, *J* = 7.5 Hz, 3H).

**$^{13}$C NMR** (101 MHz, $CD_3OD$) δ 135.70, 134.00, 130.21, 128.71, 121.77, 119.42, 117.64, 116.15, 112.34, 107.36, 61.19, 55.95, 51.22, 44.61, 36.54, 29.70, 27.02, 19.98, 15.76, 7.70.

**$^{1}$H NMR of vinpocetine alcohol:** (400 MHz, $CD_3OD$)

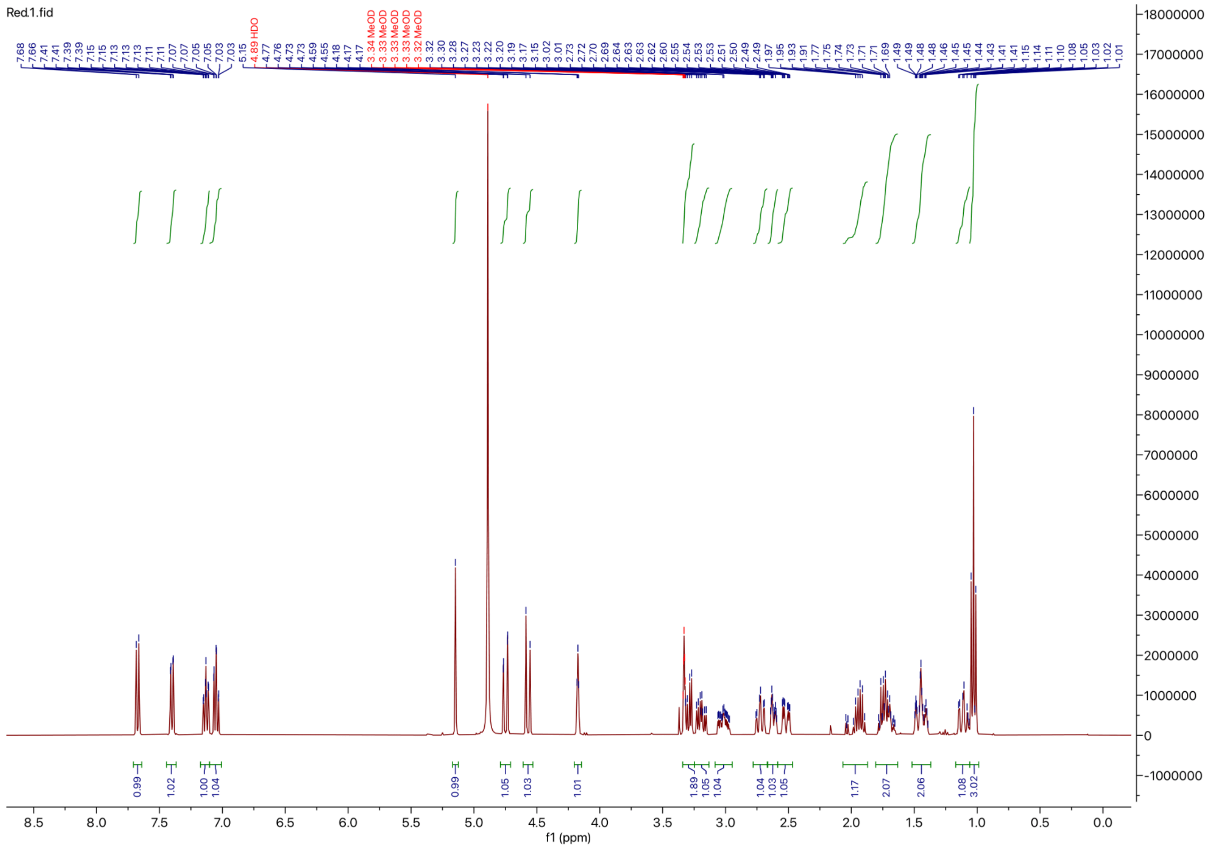

**$^{13}$C NMR of vinpocetine alcohol:** (101 MHz, $CD_3OD$)

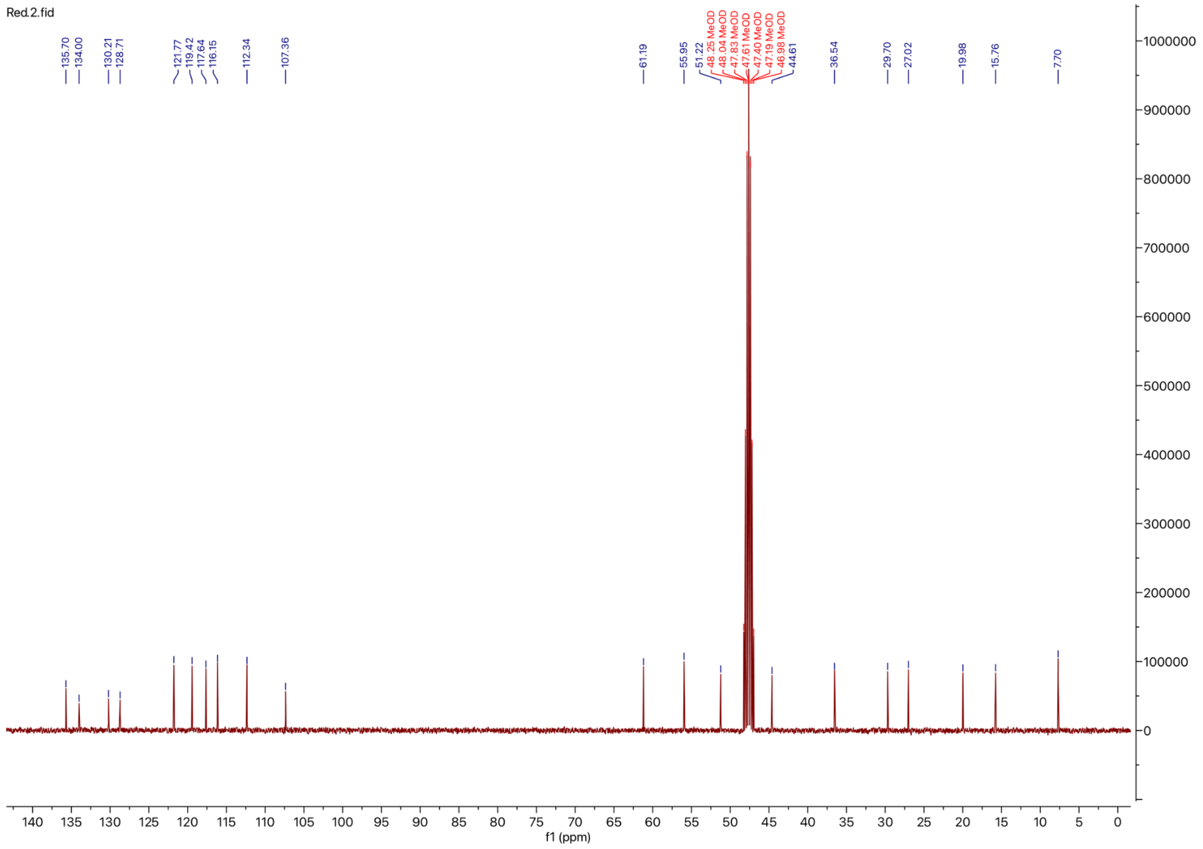

#### b. Compound 1.4 Vinpocetine Acid

**Preparation of Vinpocetine Acid (1.4)**

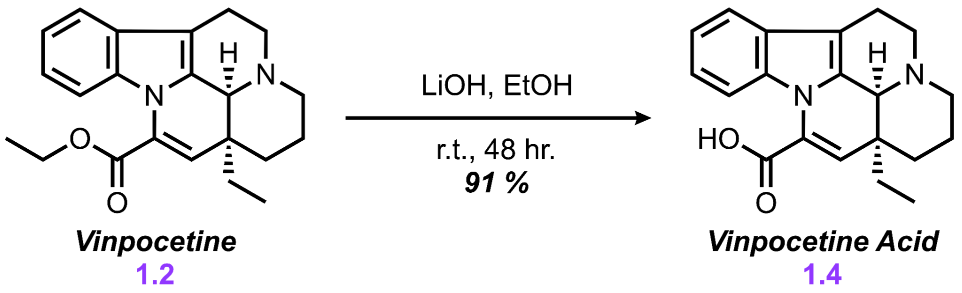


**Chemicals:**
Vinpocetine (AK Scientific, 95%): used without further purification
Lithium hydroxide (Sigma Aldrich, 97%): used without further purification
Ethyl alcohol (Acros organics, 99.5%): used without further purification

**Procedure:** To a 25 mL round bottom flask fitted with a Teflon stir bar was added Vinpocetine (350. mg, 1.0 mmol, 1.0 equiv) and lithium hydroxide (240 mg, 10 mmol, 10 equiv) and the mixture was suspended in ethanol (12 mL). The reaction was stirred at room temperature for 48 hr. after which TLC analysis indicated full hydrolysis of ester **1.2** to acid **1.4**. The resulting mixture was partially evaporated *in vacuo* and directly loaded onto a silica column. The resulting compound was eluted using a gradient of 30% to 80% ethyl alcohol in hexanes) to afford acid **1.4** (294 mg, 91%) as white solid powder.

**Characterization Data of Vinpocetine Acid 1.4**

**TLC ($R_f$):** 0.3 (95% EtOH/ 5% N-Heptane), UV active, purple-blue spot by ninhydrin stain

**LCMS (ESI-MS)**: Calculated for$[C_{20}H_{22}N_2O_2Na]^+[M+Na]^+$: 345.398; found: 345.676

**FT-IR (ATR, $cm^{-1}$)**: 3359.75, 2937.92, 2358.63, 2251.15, 1593.93, 1455.45, 1403.83, 1357.11, 1276.29, 1083.49, 1022.13, 904.33, 805.86, 726.00, 648.93, 604.22

**$^1$H NMR** (400 MHz, $CD_3OD$) δ 7.54 (dd, *J* = 8.0, 1.1 Hz, 1H), 7.51 – 7.38 (m, 1H), 7.15 – 7.00 (m, 2H), 5.66 (s, 1H), 4.28 (d, *J* = 2.7 Hz, 1H), 3.46 – 3.24 (m, 3H), 3.14 – 3.00 (m, 1H), 2.79 – 2.64 (m, 2H), 2.60 (ddd, *J* = 16.1, 5.2, 2.0 Hz, 1H), 2.04 – 1.62 (m, 3H), 1.62 – 1.53 (m, 1H), 1.46 (dt, *J* = 13.7, 3.2 Hz, 1H), 1.24 – 1.12 (m, 1H), 1.16 – 1.00 (m, 4H).

**$^{13}$C NMR** (101 MHz, $CD_3OD$) δ 170.29, 134.46, 134.01, 129.68, 128.47, 121.19, 119.37, 118.68, 117.37, 112.36, 106.80, 55.88, 51.15, 44.59, 37.01, 29.19, 27.09, 19.66, 15.79, 7.63.

**$^{1}$H NMR of vinpocetine acid:** (400 MHz, $CD_3OD$)

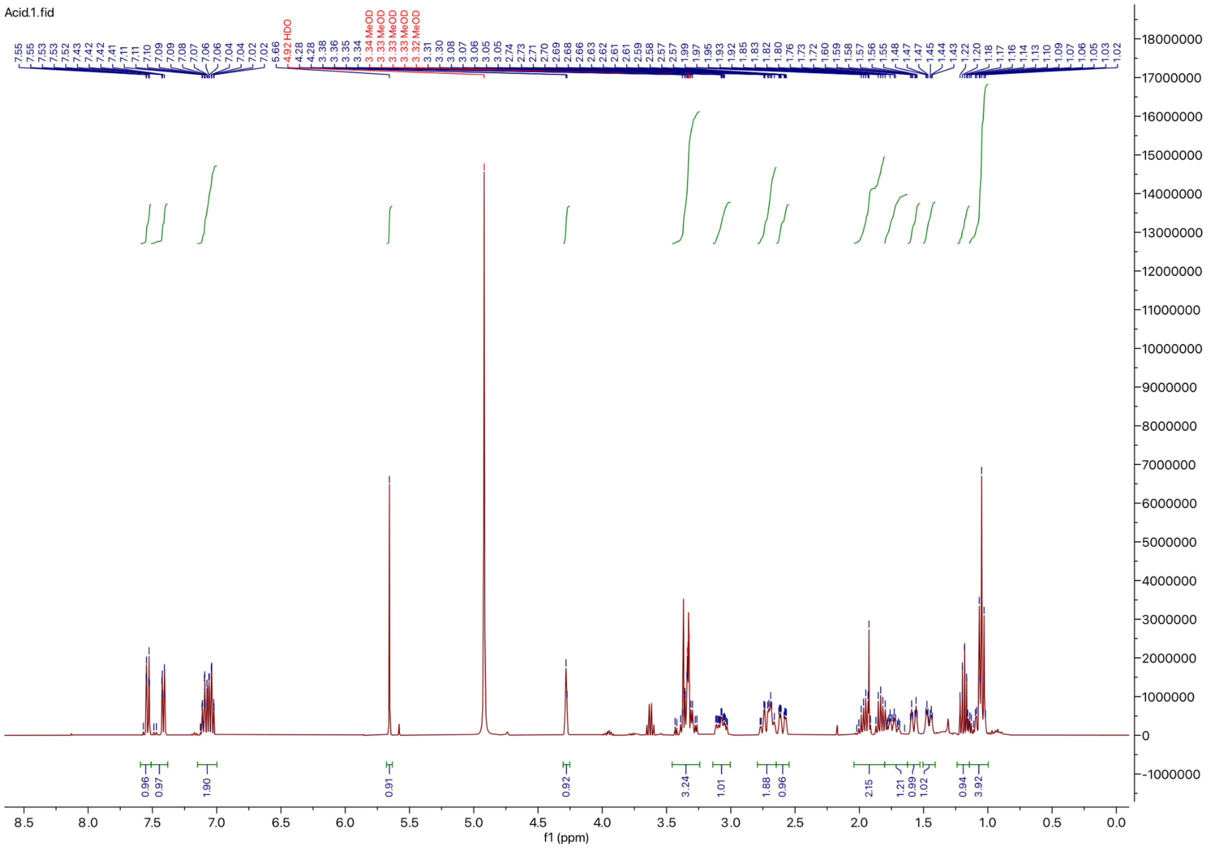

**$^{13}$C NMR of vinpocetine acid:** (101 MHz, $CD_3OD$)

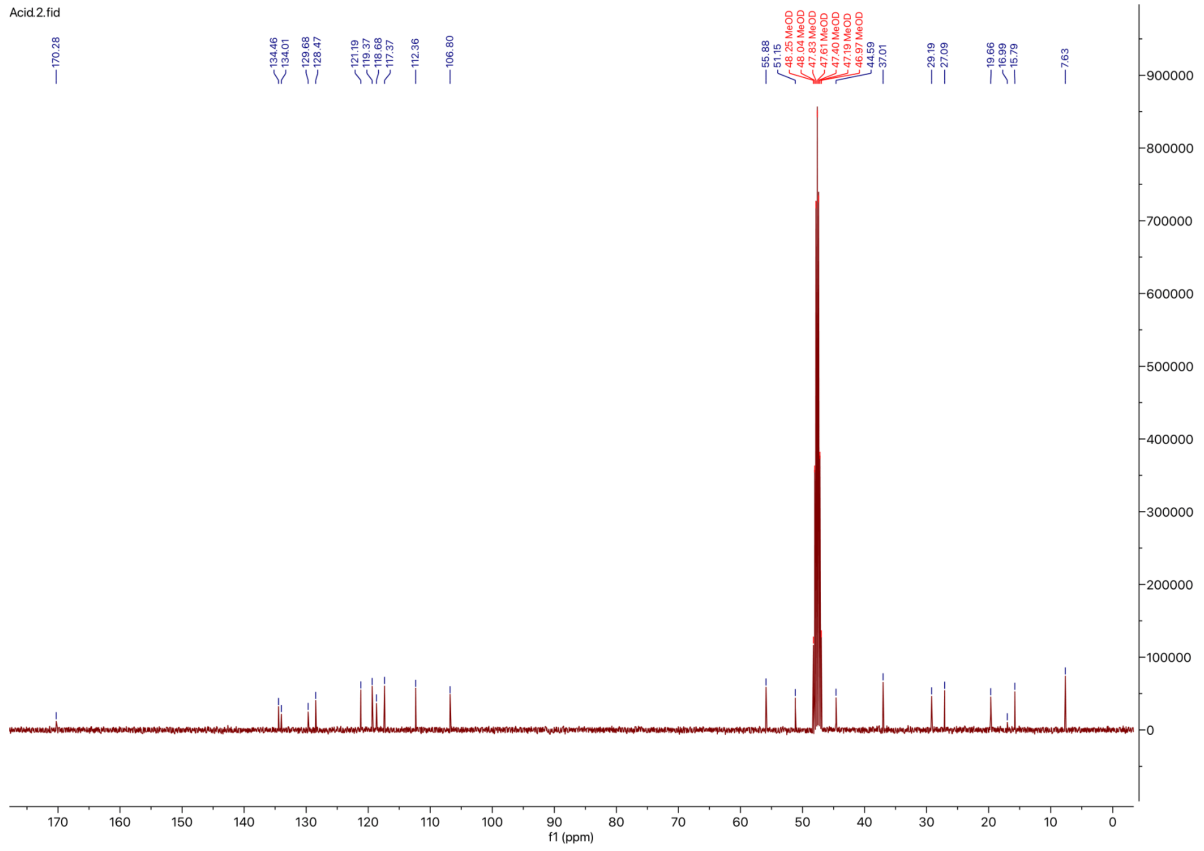

### D. Supplemental Tables

**Table 1.** Table of p-values with respect to DMSO control with and without CHIR-99021 for Wnt1-dependent luciferase reporter cell assay

| Compound (w/o CHIR-99021) | 500 µM | 250 µM | 50 µM | 25 µM |
|---|---|---|---|---|
| Vincamine **1.1** | 0.843954 | 0.359889 | 0.427374 | 0.63569 |
| Vinpocetine **1.2** | 0.011659 | 0.072788 | 0.29225 | 0.338 |
| Vinpocetine Alcohol **1.3** | 0.007157 | 0.03306 | 0.169023 | 0.209756 |
| Vinpocetine Acid **1.4** | 0.005967 | 0.014621 | 0.033624 | 0.208005 |

| Compound (w/ CHIR-99021) | 500 µM | 250 µM | 50 µM | 25 µM |
|---|---|---|---|---|
| Vincamine **1.1** | 0.831986 | 0.928956 | 0.1065 | 0.121966 |
| Vinpocetine **1.2** | 0.002258 | 0.014857 | 0.001146 | 0.000248 |
| Vinpocetine Alcohol **1.3** | 0.001419 | 0.004036 | 0.000621 | 0.001133 |
| Vinpocetine Acid **1.4** | 0.001257 | 0.001685 | 0.050948 | 0.002726 |
